# A left-to-right bias in spatial numerical associations with dots and symbols

**DOI:** 10.1101/2025.07.28.666512

**Authors:** Elena Eccher, Serge Caparos, Marco Buiatti, Manuela Piazza, Giorgio Vallortigara

**Affiliations:** CIMeC, University of Trento (Italy); Laboratoire DysCo, Université Paris 8, Saint-Denis, France; Institut Universitaire de France, Paris, France

**Keywords:** Spatial Numerical Association, SNARC effect, Numerical cognition

## Abstract

Number and space are intertwined in human and non-human cognition. A substantial body of research has shown that numerical magnitudes are mentally represented along a spatially oriented continuum, akin to a “mental number line”. Some suggested that its directionality is determined by culture and context, an idea that was recently challenged by a study that showed that an implicit association between “left” and “small” emerges not only in literate adults, but also in unschooled indigenous adult populations and preschool Western children. This finding suggests that SNAs may originate from universal innate mechanisms rather than being solely a by-product of cultural learning. However, while the study reported a strong association between “left” and “decreasing” numerosity, there was only a very weak association between “right” and “increasing” numerosity. This asymmetry was not predicted and needs further investigation to be understood. Here, we investigated the number/space association in implicit tasks in educated Western adults in greater depth by using more variable and better-controlled stimuli compared to the ones used in the previous study, and also manipulating stimulus format, using both dot patterns and symbolic numbers. Fifty-one adult participants performed a numerical comparison task within a Go-No-Go paradigm on subsequent pairs of visual stimuli (with ratios spanning from 0.75 to 0.94) that could appear on the left or on the right of a fixation point and completed two different tasks: “press when more” and “press when less”. Results revealed distinct response patterns depending on the symbolic/non-symbolic nature of the stimuli. When non-symbolic stimuli were used, a consistent association between small numerosities and the left side and large numerosities and the right side was observed. When symbolic stimuli were used, only an association between large numerosities and the right side was observed. These findings support the hypothesis that SNAs may reflect a biological predisposition associated with brain asymmetry and that task demands may interact with the underlying hemispheric specialisations.

## INTRODUCTION

Animals constantly rely on information relative to the numerosity of sets for survival, whether hunting, choosing mates or avoiding predators, and this ability is widespread in the animal kingdom, from invertebrates to humans (Bortot et al., 2021; Butterworth et al., 2018; Giurfa, 2019; Nieder, 2019; Vallortigara, 2018). The Approximate Number System (ANS), or *Number Sense*, has been identified as the putative system that enables individuals to perceive, estimate and internally manipulate numerical quantities (Dehaene, 2011). This “Core System” for numbers would be rooted in our evolutionary history and available since birth (Di Giorgio et al., 2019; Lorenzi et al., 2025; Spelke, 2000; Vallortigara, 2021). One of the main features of the ANS is that, similarly to other perceptual abilities, it obeys Weber’s law, suggesting that it works solely on a ratio-dependent limit.

At the neural level, the ANS has been linked to cortical networks centered on the intraparietal sulcus (IPS) and extending to other regions in both the occipito-temporal and frontal cortices in humans (Piazza & Eger, 2016) and in pallial areas in the avian (Kobylkov et al., 2023; Nieder, 2021) and zebrafish (Luu et al., 2024) brains, where number-selective neurons, displaying log-gaussian tuning functions, have been described. In humans, functional neuroimaging has repeatedly shown that the IPS is recruited during numerosity processing tasks, such as estimating or comparing the number of visual objects (Dehaene et al., 2003; Duricy et al., 2025), that its activity is modulated by numerical ratio in adaptation paradigms (Piazza et al., 2004) and predicts subjects’ behavioural numerosity estimation precision (Eger et al., 2009; Lasne et al., 2019).

This ANS system was also suggested both to play a foundational role in the acquisition of symbolic numeracy (Piazza, 2010; Halberda et al., 2008; Decarli et al., 2023), but also to be substantially reshaped during learning, in particular in the left hemisphere (Piazza et al., 2007; Verguts & Fias, 2004). In the following, we distinguish between *numerosity*, referring to the perception of *non-symbolic* quantities through sets of objects (e.g., arrays of dots), and *numbers*, referring to culturally learned s*ymbolic* representations such as Arabic digits or number words. Developmental studies show that non-symbolic numerosity processing in both infants and preschoolers is associated with a right lateralized IPS activity (Cantlon et al., 2006; Hyde et al., 2010; Izard et al., 2008), while this activity extends towards the left IPS as children get acquainted with symbolic numbers (Aulet et al., 2025; Emerson & Cantlon, 2015; Rivera et al., 2005; Vogel et al., 2015). It therefore appears that, as language and symbolic number systems develop, the early right-lateralized response to numerosities evolves into a more bilateral pattern, reflecting the growing role of the left hemisphere in symbolic number processing (Ansari & Dhital, 2006). When directly contrasting the activity evoked by non-symbolic *vs*. symbolic quantities in adults, a different pattern of lateralisation is evident, with a higher activation of the left hemisphere in response to digits and number-words and a more right-lateralised one in response to a non-symbolic pattern of dots (Damarla & Just, 2013; Holloway et al., 2010; Piazza et al., 2007; Sokolowski et al., 2021; Zang et al., 2024).

Another lateralized aspect of both numerosity and number processing is associated with the phenomenon of Spatial-Numerical Associations (SNA), in which numerical quantities are mapped onto spatial dimensions. One example is provided by the *Spatial-Numerical Association of Response Codes* (SNARC, (Dehaene et al., 1993)), where subjects are faster to complete a parity judgment task when presented with numerical symbols with their left hand in response to small numbers and with their right hand in response to large numbers. On the contrary Iranian participants showed either the opposite or no effect at all (Dehaene et al., 1993). The authors interpreted these results as evidence that numbers are spatially organized on a so-called Mental Number Line (MNL, (Galton, 1880; Restle, 1970), and that its orientation is determined by the direction of reading/writing habits. In following years several studies seemed to confirm this cultural explanation for the direction of the SNA, especially stressing that the directionality of the cultural habits of reading and writing might account for the observed effects. For example, left-to-right readers often show a left-to-right SNARC effect, while right-to-left readers tend to exhibit reversed or absent effects (Fischer et al., 2009; Fischer & Rottmann, 2005; Göbel et al., 2011; Nuerk et al., 2004; Zebian, 2005; Zohar-Shai et al., 2017). This idea was also supported by the observation that an indigenous population with no writing system showed no consistent spatial-numerical mapping when asked to order numerosities on a line ((Pitt et al., 2021).

However, a concurrent line of research proposes that the SNA might not be *determined* by cultural factors, but rather it can be *modulated* by them. This position is determined by the fact that some forms of SNA are present in non-human animals species (that clearly lack knowledge of written language or culture-based counting or ordering practices), such as monkeys (Adachi, 2014; Brannon & Terrace, 1998; Rugani et al., 2024), birds (Rugani et al., 2010, 2015, 2020), zebrafish (Potrich et al., 2026), and even bees (Giurfa et al., 2022). Similarly, preverbal infants and neonates also exhibit a consistent left-to-right mapping of numerical quantity, suggesting that, even in humans space-number mappings might start well before the acquisition of spatialized cultural practices (Bulf et al., 2016; De Hevia et al., 2014; de Hevia, Veggiotti, et al., 2017; Di Giorgio et al., 2019). It has been suggested that this spontaneous left-to-right spatial-numerical mapping could be the result of lateralized brain functions. Vallortigara (2018) hypothesised that SNA may stem from hemispheric specialisation for approach and withdrawal behaviours, with the left hemisphere processing positive-valence (approach) stimuli and the right hemisphere processing negative-valence (withdrawal) stimuli (Davidson, 2004). In this view, increasing numerosities, typically associated with positive valence in the real world (e.g. associated with rewards), would bias attention to the right hemispace, while decreasing numerosities, linked to negative valence, would shift attention to the left hemispace (Vallortigara, 2018). This growing body of research has led to the hypothesis that, in human adults, SNA may arise from a combination of biological and culturally-mediated mechanisms (Eccher et al., 2026; Eccher & Vallortigara, 2026; Guida et al., 2018).

Thus, while SNA might be universal in its origin, cultural context, individual experiences and task demands can modulate its specific appearance. In particular, the demands of the tasks used to study SNA might influence the possibility of eliciting different aspects of it, either in favour of the biological predisposition or in favour of the cultural acquisitions. Evidence from implicit SNA tasks in human adults, similar to those found in non-human animals and preverbal infants, suggests a role for biologically grounded spatial-numerical associations, whereas more explicit tasks may rely more on culturally acquired spatial-numerical mappings, shaped for example by reading and writing habits (Eccher et al., 2025). To test this, in our previous work we compared performance on two tasks across three groups with different literacy exposure: Italian adults (high literacy), Italian preschoolers (minimal literacy), and Himba adults (an indigenous population from northern Namibia with an oral culture and no writing system). In the explicit card-ordering task, where participants were asked to order cards in space on the bases of their numerical values, only Italian adults showed a consistent left-to-right SNA aligned with their cultural reading direction, while preschoolers and Himba participants showed no directional bias, despite favouring lateral linear arrangements (over vertical, obliquus, circular, or random ones). In contrast, in the implicit Go/No-Go task, where the spatial position of the stimuli was determined by the experimenter, and completely task irrelevant, all three groups, including the Himba, exhibited similar left-to-right oriented SNAs, suggesting that implicit number-space associations may rely on biologically rooted mechanisms rather than on acquired cultural practices (Eccher et al., 2025).

However, while this previous study did indeed suggest that such a clear dissociation between explicit and implicit form of SNA might reflect the different contributions of biological predispositions and cultural acquisitions, an interesting and unexpected asymmetrical Congruency Effect during the implicit task was observed: participants across groups performed faster and more accurately when numerically smaller stimuli appeared in the left visual field in the decreasing task, while no significant effect was observed for larger stimuli appearing in the right visual field in the increasing task. This pattern held across differences in participants’ age and cultural background. We hypothesized that this asymmetry may stem from the non-symbolic nature of the stimuli we used and related it to the right-hemispheric superiority for numerosity. According to this interpretation, the right hemisphere may have been more strongly engaged during non-symbolic numerosity processing, facilitating performance in the left visual field during the decreasing task and weakening the effect in the right visual field during the increasing one. However, if this brain asymmetric effect was the only factor determining subjects’ behaviour, we should have observed a fully reversed Congruency Effect in the increasing task, instead of a null effect in this condition. Thus, the fact that we did not observe a facilitation for any stimulus presented in the left visual field, but only for small numerosities, suggests that the results cannot be solely explained by the hemispheric superiority for numerosity. We thus hypothesized that the asymmetry of the effect could also be partially explained by an inherent difference in difficulty across the increasing and decreasing tasks. In the literature, there is indeed evidence of a broader cognitive advantage for processing of increasing *vs*. decreasing quantities (see also for example the so-called “addition advantage”, Barth et al., 2006, 2008; Gilmore & Spelke, 2008; Kamii et al., 2001). Given the adaptive advantages of rapidly detecting increases in critical resources, the human brain may be evolutionarily biased toward rising numerosities. For example, it is well known that since early infancy we have an advantage for processing increasing *vs*. decreasing ordinal sequences (de Hevia, Addabbo, et al., 2017; Macchi Cassia et al., 2012), and asymmetry emerges also in adults when comparing task instructions in SNARC paradigms (Patro et al., 2015; Patro & Haman, 2011; Shaki et al., 2012). It is therefore possible that facilitating mechanisms for processing increasing changes in numerosity (Ben-Meir et al., 2012; Ganor-Stern, 2015; Müller & Schwarz, 2008) led to faster responses for the increasing task in our first study, which might have masked the more subtle SNA effect.

In sum, it is possible that in our original study, SNA-related mechanisms may have been partially obscured in the increasing task condition, either due to differential hemispheric recruitment or to task demands or both. To test these hypotheses, we designed a new study with a modified version of our original task. While retaining the core design, we introduced two critical manipulations. First, we varied the numerosity presentation modality by including not only non-symbolic (dots) stimuli, but also including symbolic numbers (digits and number words). This allowed us to test the impact of the stimulus format on the SNA. We expected that the asymmetrical pattern observed in Eccher et al. (2025) would reverse when the task involved numerical symbols, predicting in such cases a stronger SNA effect in the increasing condition, reflecting the well-known left-hemispheric superiority for symbolic number processing. Second, we increased the difficulty of the numerical comparison task, hypothesizing that more cognitively demanding tasks would reduce the impact of the facilitatory effect of the increasing task, and thus enhance the likelihood of detecting bilateral SNA patterns, at least for non-symbolic stimuli.

## METHODS

The study adhered to the principles outlined in the Declaration of Helsinki. All participants gave written informed consent after receiving a comprehensive explanation of the study’s purpose and procedures. A small monetary reward was provided for their participation.

In the same session, we conducted three separate experiments, each utilizing a distinct modality of stimulus presentation. The task was modeled after the Numerosity Comparison Task described in our earlier study (Eccher et al., 2025), with minor procedural variations (see below).

### Participants

Fifty-one healthy adults from the Trentino region in Italy (40 females; mean age = 24.63 ± 5.25 years) were recruited through a social media group affiliated with the University of Trento. All participants completed the Dots experiment, while thirty-nine of them also took part in the Digits and Words experiments.

### Stimuli

We included one non-symbolic and two symbolic stimulus formats, each used in a distinct task. In all conditions, the reference stimulus was 16 and the test stimuli were 12, 13, 14, 15, 17, 18, 19 and 20. While the reference stimulus was always presented at the center of the screen, the test stimuli were displayed 15° to the left or right of the central fixation point.

#### Dots

The non-symbolic stimuli consisted of randomly arranged black squares of a constant size (1.3 × 1.3 degrees of visual angle) presented against a white background of 17 × 17 degrees, similarly to the paradigm used in Eccher et al., 2025 (Fig 1A).

**Figure 1.**
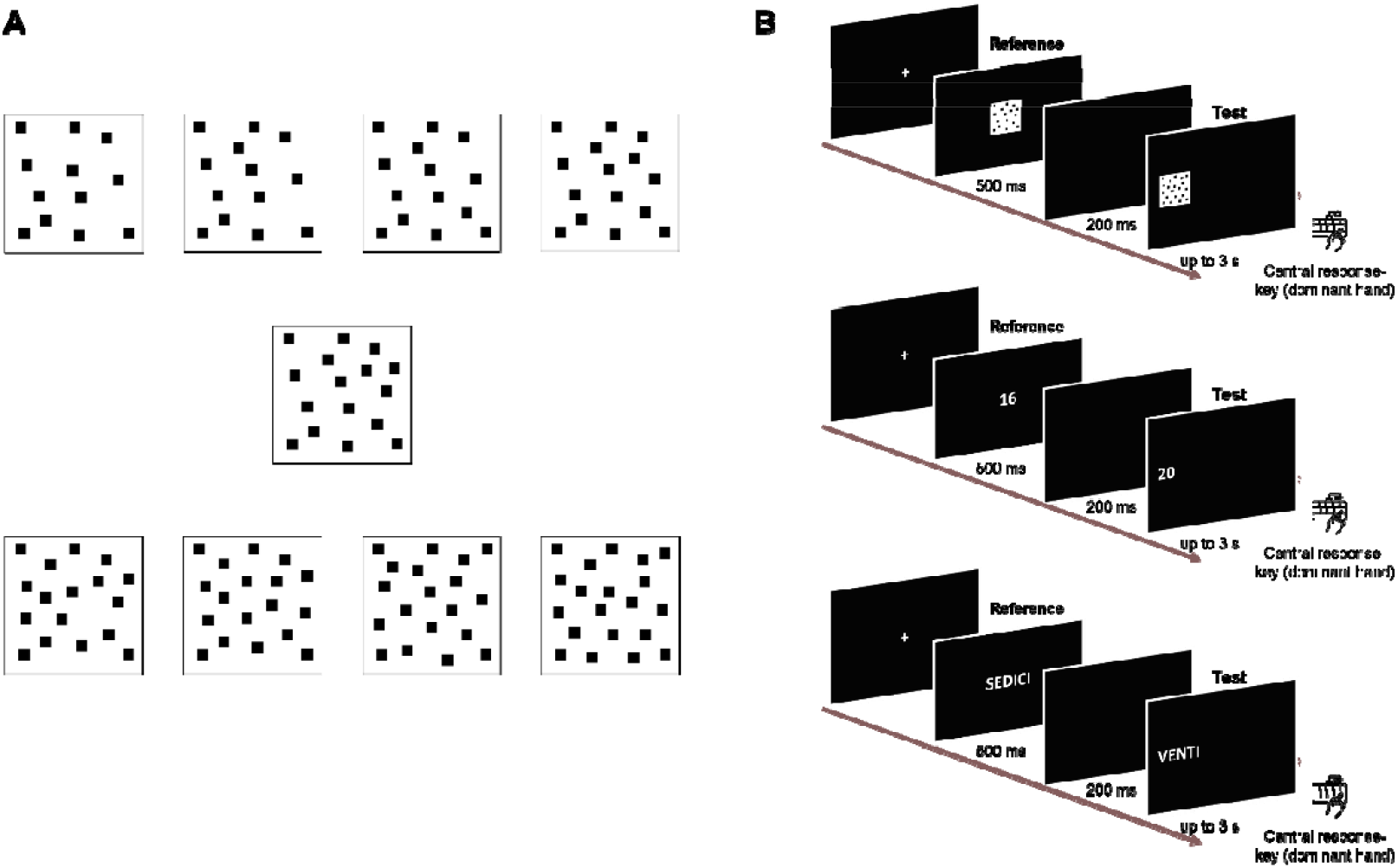
Schematic representation of paradigm and stimuli. Panel A) Stimuli used in the Dots experiment. On the top, from left to right, numerosities from 12 to 15; in the center, reference numerosity 16; at the bottom, from left to right, numerosities from 17 to 20. Panel B) Schematic representation of the paradigm design, from top to bottom an example trial with the three different types of stimuli used (Dots arrays, Arabic digits and Number Words).

#### Digits

Symbolic stimuli in the digit format were Arabic numbers shown in white Arial font (85 pt) on a black background.

#### Words

Symbolic stimuli in the word format were displayed as capitalized letters in white Arial font (57 pt) against a black background.

### Procedure

Participants were tested in a dimly lit lab at the **anonymised lab center**. Stimuli were shown on a computer monitor, positioned about 50 cm from the participant. The full session lasted around 40 minutes, and participants were allowed to take breaks between each task, if needed. For each stimulus modality (Dots, Digits, Words) subjects performed two tasks: Increasing and Decreasing (see below). Each stimulus modality was tested independently, meaning that no task involved a comparison between two different stimulus modalities. To avoid priming effects, the Dots task was always performed first. The symbolic tasks (Digits and Words) followed, with their order counterbalanced among participants. The order of increasing/decreasing tasks was also counterbalanced across subjects. The reference stimulus was fixed at “16” in each trial. Each trial began with a 1-second central fixation cross, followed by the reference stimulus for 500 ms. After a 200 ms blank screen, the test stimulus appeared on either the left or right side. It remained visible until the participant responded or until 3 seconds elapsed (Fig 1B). Participants were instructed to press a central key as quickly as possible, using their dominant hand, only if the test stimulus had fewer items than the reference (Decreasing Task) or more items (Increasing Task). Feedback was given throughout the entire experiment, using a green happy face for correct answers and a red sad face for incorrect ones, both displayed for 1 second. Each condition (Increasing and Decreasing) included 50 trials: 32 Go trials (8 per numerosity pair) and 18 No-Go trials. This unbalance across conditions mirrored the one present in the original experiment (Eccher et al, 2025).

### Statistical analyses

All data analyses and graphical outputs were carried out using R (version 4.1.3) within the RStudio environment. An alpha level of .05 was adopted for determining statistical significance. For effect size reporting, we computed the effect size C for Wilcoxon tests (i.e., 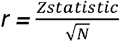). For ANOVAs, partial eta squared served as the measure of effect size.

Reaction times (RTs) were gathered in Go trials under both congruent and incongruent conditions for each participant. As in Eccher et al., 2025 we defined the Congruency condition with respect to the left-to-right space to number mapping: targets that were smaller than the reference and appeared on the left, as well as those that were larger and appeared on the right, were classified as congruent. Conversely, smaller targets on the right and larger targets on the left were considered incongruent. Prior to the statistical analysis, outlier trials were removed for each experimental condition (Dots, Digits, Words). In order to clean the data from outliers, trials were excluded if RTs fell below the first quartile minus 1.5 times the interquartile range (IQR) or exceeded the third quartile plus 1.5 times the IQR.

First, we examined the influence of numerical distance between the reference and test stimuli on reaction times using a 3 (Format) × 4 (Numerical Distance) repeated-measure ANOVA. Then, we conducted a repeated measures ANOVA with three within-subject factors: Format (Dots, Digits, Words), Task (Increasing, Decreasing), and Congruency condition (Congruent, Incongruent). In cases where Mauchly’s test indicated a violation of the sphericity assumption (p ≤ .05), the Greenhouse-Geisser correction was applied to the affected factors, and adjusted results are reported accordingly. Wilcoxon signed-rank tests were used for post hoc comparisons between congruent and incongruent conditions within each Task and Stimulus format.

To replicate the analytical approach used in Eccher et al, 2025, we also calculated the RTs Congruency Effect for each task using the 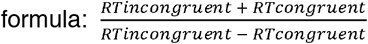. We then performed a 3 (Format) × 2 (Task) repeated measures ANOVA to examine the effects of experimental condition and task instruction. The Congruency Effect was further evaluated against zero using a two-tailed Wilcoxon signed-rank test. It is important to note that, due to incomplete participation across all experimental conditions, the repeated measures ANOVAs were limited to the 39 participants who completed every experiment. Instead, post-hoc analyses were conducted using the full available dataset for each condition, as indicated by the degrees of freedom in the results tables.

Finally, to be able to better discuss the results in comparison to our previous findings, a 2×2 mixed ANOVA, with study (Eccher e al., 2025 vs current study) as between factor, and Task as within factor was carried out to compare the reaction times from the present work with the data from Eccher et al., 2025 (see Supplementary Materials).

## Results

Diversely from what was done in Eccher et al., 2025 in the present study the analyses are based solely on reaction times (RTs) rather than on the Inverse Efficiency Score (IES). The reason for this choice is that the current sample consisted exclusively of adult participants, whose overall accuracy was very high (mean accuracy = 94.7%, SEM = 0.3%). In contrast, in Eccher et al., 2025 the sample included children and Himba participants, who showed a substantially higher number of errors. In that context, a normalised score, namely the IES, was needed to allow a fair comparison across populations with different accuracy levels.

First of all, as a form of sanity check to ensure that the task evoked a representation of magnitude, we investigated the distance effect as its main signature, and its relation with the format with which the numerical stimuli were presented (as dots, as digits, or as number words). To examine these effects, we performed a 3 × 4 repeated-measures ANOVA on RTs with Numerical Distance and Format as within-subject factors. The analysis revealed a significant main effect of Numerical Distance, F_(1.47, 54.32)_ = 93.47, p < .001, η^2^p = .716, and a significant main effect of Format, F_(1.44, 53.39)_ = 65.27, p < .001, η^2^p = .638. Consistent with the typical distance effect, RTs were significantly shorter when the numerical distance between the reference and the test stimulus was larger. In addition, a main effect of Format indicated that responses were significantly slower for word stimuli and faster for Digit stimuli. Numerical Distance also interacted with Format (F_(1.69, 62.39)_ = 16.146, p < .001, η_p_ ^2^ = 0.304, see Figure S1), indicating that even if distance modulated RTs in all conditions, this effect was stronger for the non-symbolic format (Table 1, Fig. S1).

**Table 1.** Means and Standard Errors of the mean for each numerical distance in each format.

| **Reaction Times per Numerical Distance** |  |  |  |  |
| --- | --- | --- | --- | --- |
| *Format* | *Numerical Distance* | n | Mean (s) | SEM |
| Dots | 1 | 51 | 0.733 | 0.028 |
|  | 2 | 51 | 0.605 | 0.015 |
|  | 3 | 51 | 0.554 | 0.011 |
|  | 4 | 51 | 0.541 | 0.01 |
| Digits | 1 | 39 | 0.539 | 0.009 |
|  | 2 | 39 | 0.504 | 0.008 |
|  | 3 | 39 | 0.516 | 0.007 |
|  | 4 | 39 | 0.498 | 0.009 |
| Words | 1 | 39 | 0.702 | 0.015 |
|  | 2 | 39 | 0.644 | 0.012 |
|  | 3 | 39 | 0.624 | 0.012 |
|  | 4 | 39 | 0.613 | 0.01 |

Given the limited number of trials per subject for each test numerosity, and our primary interest in examining the overall direction of numerosity changes in relation to Task, we then collapsed reaction times across the different levels of test numerosity within each of the two Tasks. This allowed us to examine the effect of congruency between the direction of numerosity change and format.

A 3×2×2 ANOVA was conducted to investigate the effect of Format, Task and Congruency condition on RTs (Table 2). The analysis revealed a main effect of Format and a main effect of Congruency condition. Indeed, in line with the previous analysis, there were significantly shorter RTs in response to Digits (mean = 0.501 s, SEM = 0.004 s) compared to Dots (mean = 0.560 s, SEM = 0.007 s) and Words (mean = 0.629 s, SEM = 0.006 s) and overall shorter RTs were found in the congruent condition (mean = 0.557 s, SEM = 0.006 s) compared to the incongruent condition (mean = 0.569 s, SEM = 0.006 s).

**Table 2.**
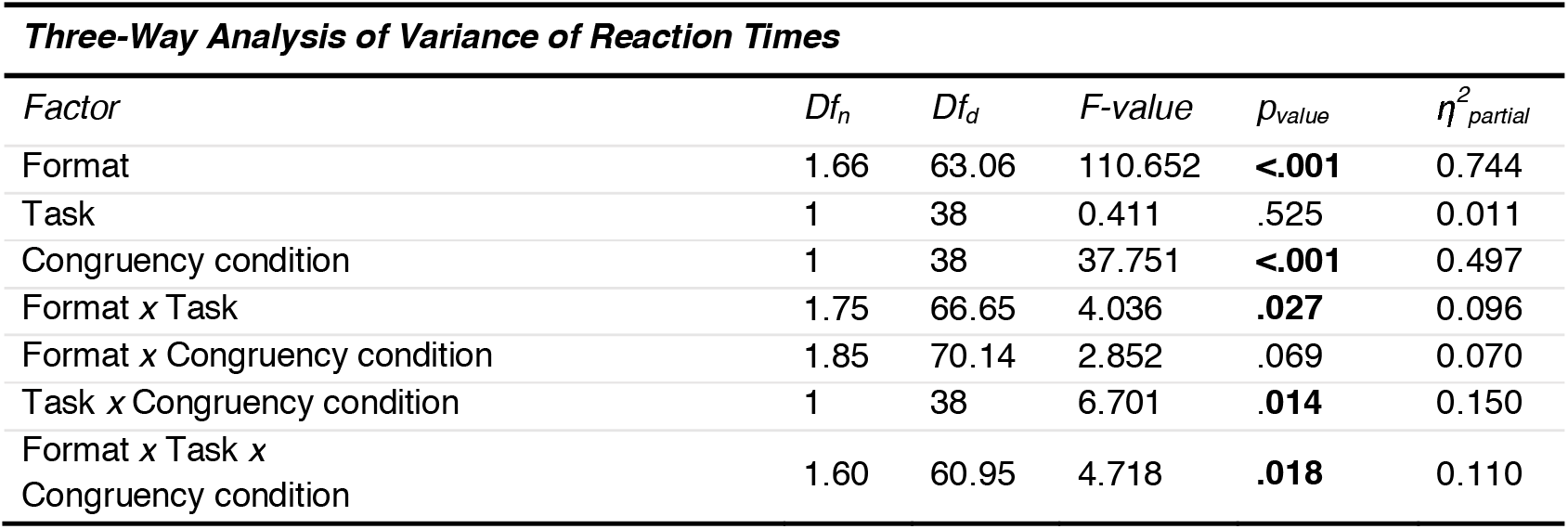
Three-Way Analysis of Variance of Reaction Times for each Format by Task and Congruency condition. Greenhouse-Geisser correction for the violation of the sphericity assumption has been applied. Significant p_value_ for .05 significance level in bold.

| Factor | $Df_n$ | $Df_d$ | F-value | $p_{value}$ | $\eta^2_{partial}$ |
| --- | --- | --- | --- | --- | --- |
| Format | 1.66 | 63.06 | 110.652 | **<.001** | 0.744 |
| Task | 1 | 38 | 0.411 | .525 | 0.011 |
| Congruency condition | 1 | 38 | 37.751 | **<.001** | 0.497 |
| Format x Task | 1.75 | 66.65 | 4.036 | **.027** | 0.096 |
| Format x Congruency condition | 1.85 | 70.14 | 2.852 | .069 | 0.070 |
| Task x Congruency condition | 1 | 38 | 6.701 | **.014** | 0.150 |
| Format x Task x Congruency condition | 1.60 | 60.95 | 4.718 | **.018** | 0.110 |

Given the significant triple interaction of Format *Task*Congruency condition, we further analysed the data by running pairwise comparisons between congruent and incongruent conditions for each Format and Task (Table 3). The analysis revealed that in the non-symbolic condition, both in the increasing and decreasing tasks, there were significantly shorter RTs in response to the congruent condition compared to the incongruent condition, while in both symbolic conditions, a significant difference was found only in the increasing task (Figure 2).

**Table 3.**
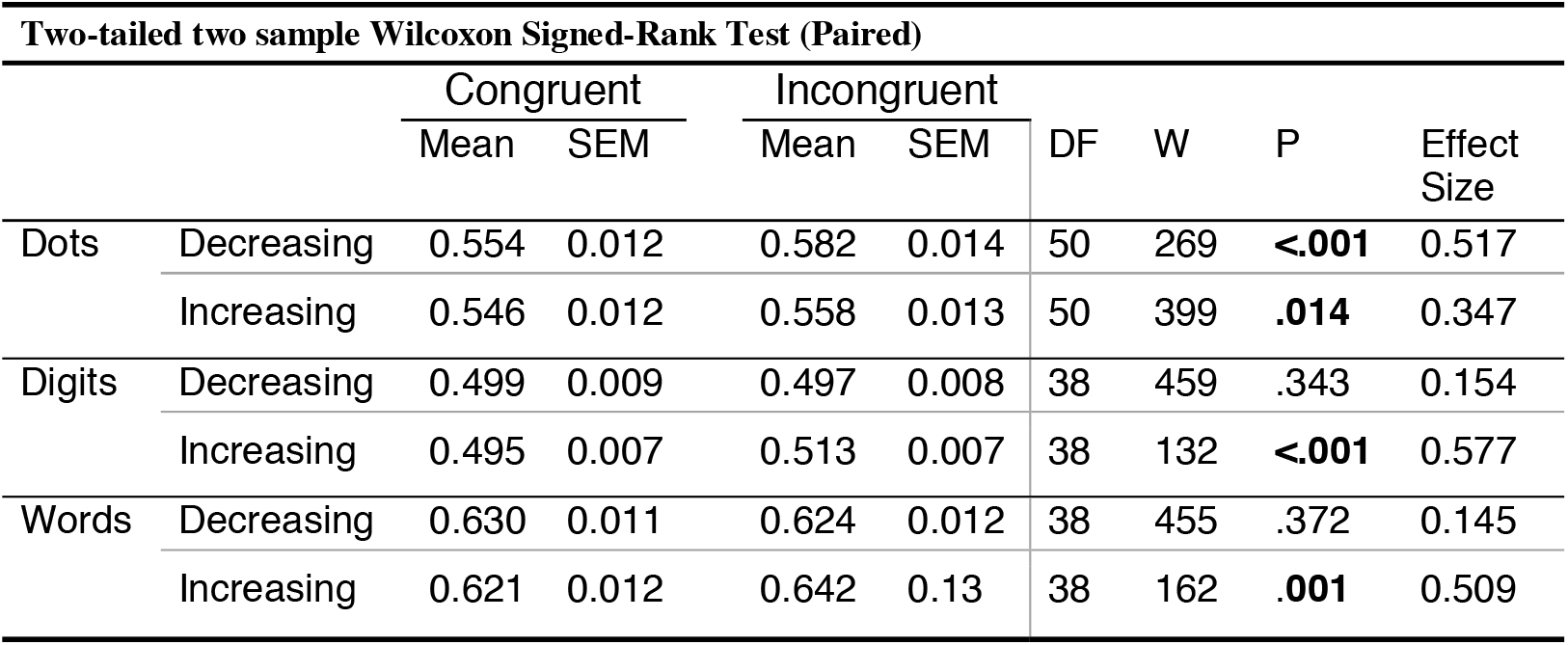
Reaction Times comparison between the two Congruency conditions for Task for each Format. Significant p_value_ for .05 significance level in bold.

| Two-tailed two sample Wilcoxon Signed-Rank Test (Paired) |  |  |  |  |  |  |  |  |  |
| --- | --- | --- | --- | --- | --- | --- | --- | --- | --- |
|  |  | Congruent |  | Incongruent |  | DF | W | P | Effect Size |
|  |  | Mean | SEM | Mean | SEM |  |  |  |  |
| Dots | Decreasing | 0.554 | 0.012 | 0.582 | 0.014 | 50 | 269 | **<.001** | 0.517 |
|  | Increasing | 0.546 | 0.012 | 0.558 | 0.013 | 50 | 399 | **.014** | 0.347 |
| Digits | Decreasing | 0.499 | 0.009 | 0.497 | 0.008 | 38 | 459 | .343 | 0.154 |
|  | Increasing | 0.495 | 0.007 | 0.513 | 0.007 | 38 | 132 | **<.001** | 0.577 |
| Words | Decreasing | 0.630 | 0.011 | 0.624 | 0.012 | 38 | 455 | .372 | 0.145 |
|  | Increasing | 0.621 | 0.012 | 0.642 | 0.13 | 38 | 162 | **.001** | 0.509 |

**Figure 2.**
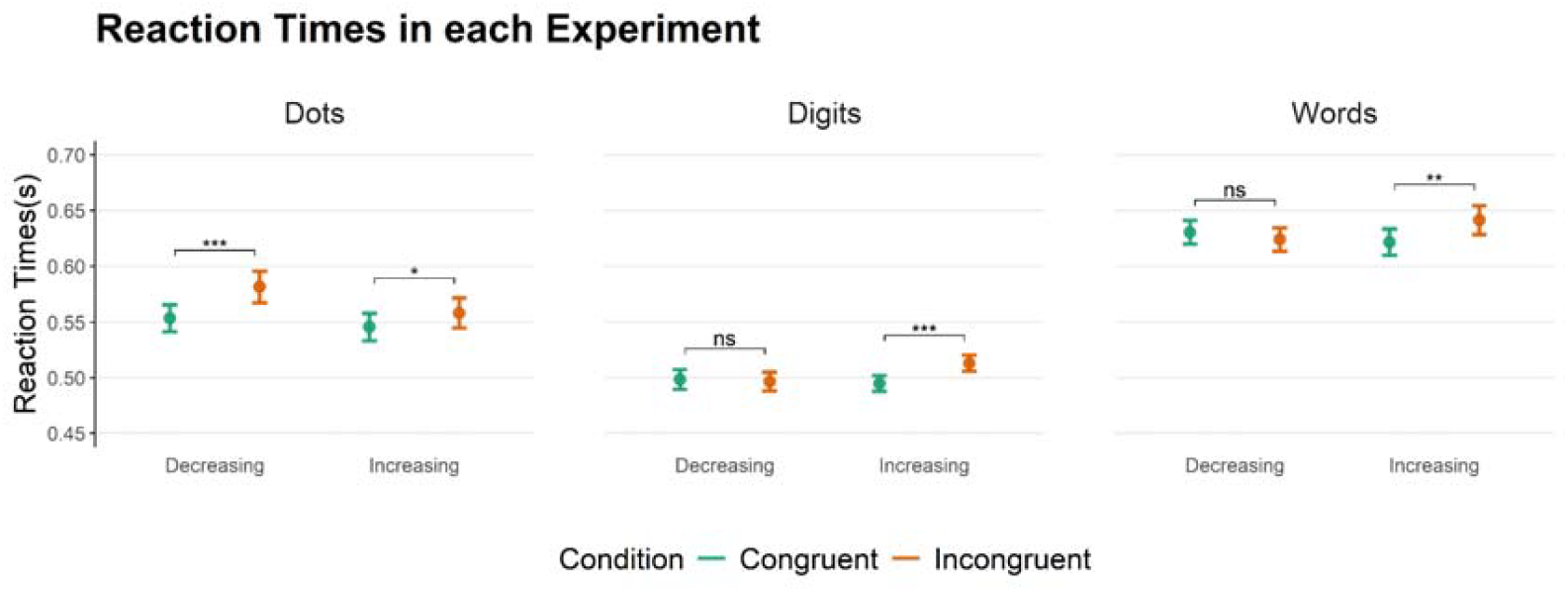
Reaction Times were measured for each Congruency Condition for each Format and Task. In the graph, mean values and standard errors of the mean are reported. Two-tailed two-sample paired Wilcoxon signed-rank test statistic, significance levels are defined as follows: * = pvalue <0.05, ** = pvalue <0.01, *** = pvalue <0.001

Similar to the Eccher et al., 2025, we also directly tested the RTs Congruency Effect (see Methods for details). To this end, a 3 × 2 repeated-measures ANOVA was conducted on the measured Congruency Effect, with Format and Task as within-subject factors. The analysis revealed no main effect of Format (F_(2, 76)_ = 2.850, p_value_ = .064, η^2^_p_ = 0.070), a significant main effect of Task (F_(1,38)_ = 6.069, p_value_ = .018, η^2^_p_ = 0.138), and interaction (F_(1.66, 63.27)_ = 5.046, p_value_ = .013, η_p_^2^ = 0.117). An overall higher Congruency Effect was found in the increasing task (mean = 0.014 s, SEM = 0.003) compared to the decreasing task (mean = 0.007 s, SEM = 0.003). No significant difference was found in Congruency Effect between decreasing and increasing tasks in response to Dots, while a significant difference was found between increasing and decreasing tasks both in response to Digits and Words (Table 4).

**Table 4.** Difference in the Congruency Effect measured from RTs for each Format and Task, significant p_value_ for .05 significance level in bold.

| Two-tailed two sample Wilcoxon Signed-Rank Test (Paired) |  |  |  |  |  |  |  |  |
| --- | --- | --- | --- | --- | --- | --- | --- | --- |
|  | Decreasing |  | Increasing |  | DF | W | P | Effect Size |
|  | Mean | SEM | Mean | SEM |  |  |  |  |
| Dots | 0.024 | 0.006 | 0.010 | 0.005 | 51 | 822 | 0.137 | 0.209 |
| Digits | -0.002 | 0.004 | 0.018 | 0.004 | 39 | 146 | <.001 | 0.545 |
| Words | -0.005 | 0.005 | 0.016 | 0.004 | 39 | 218 | 0.016 | 0.384 |

Comparison against chance level revealed a significant Congruency Effect for both the decreasing and increasing Task in the Dots experiment. On the other hand, a significant Congruency Effect for the increasing Task only was found in the Digits and Words experiment (Table 5, Figure 3).

**Table 5.** Statistical analysis against chance level for Congruency Effect for Format and Task, significant pvalue for .05 significance level in bold.

| *One-sample Wilcoxon signed rank test against chance level ($\mu = 0$)\** |  |  |  |  |  |
| --- | --- | --- | --- | --- | --- |
| *Test statistic* |  |  |  |  |  |
| *Format* | *Task* | *Df* | *t-value* | *p<sub>value</sub>* | *Effect size* |
| *Dots* | *Decreasing* | 50 | 1062 | **<.001** | 0.524 |
|  | *Increasing* | 50 | 910 | **.021** | 0.324 |
| *Digits* | *Decreasing* | 38 | 324 | .365 | 0.147 |
|  | *Increasing* | 38 | 647 | **<.001** | 0.574 |
| *Words* | *Decreasing* | 36 | 305 | .241 | 0.159 |
|  | *Increasing* | 36 | 625 | **<.001** | 0.530 |

**Figure 3.**
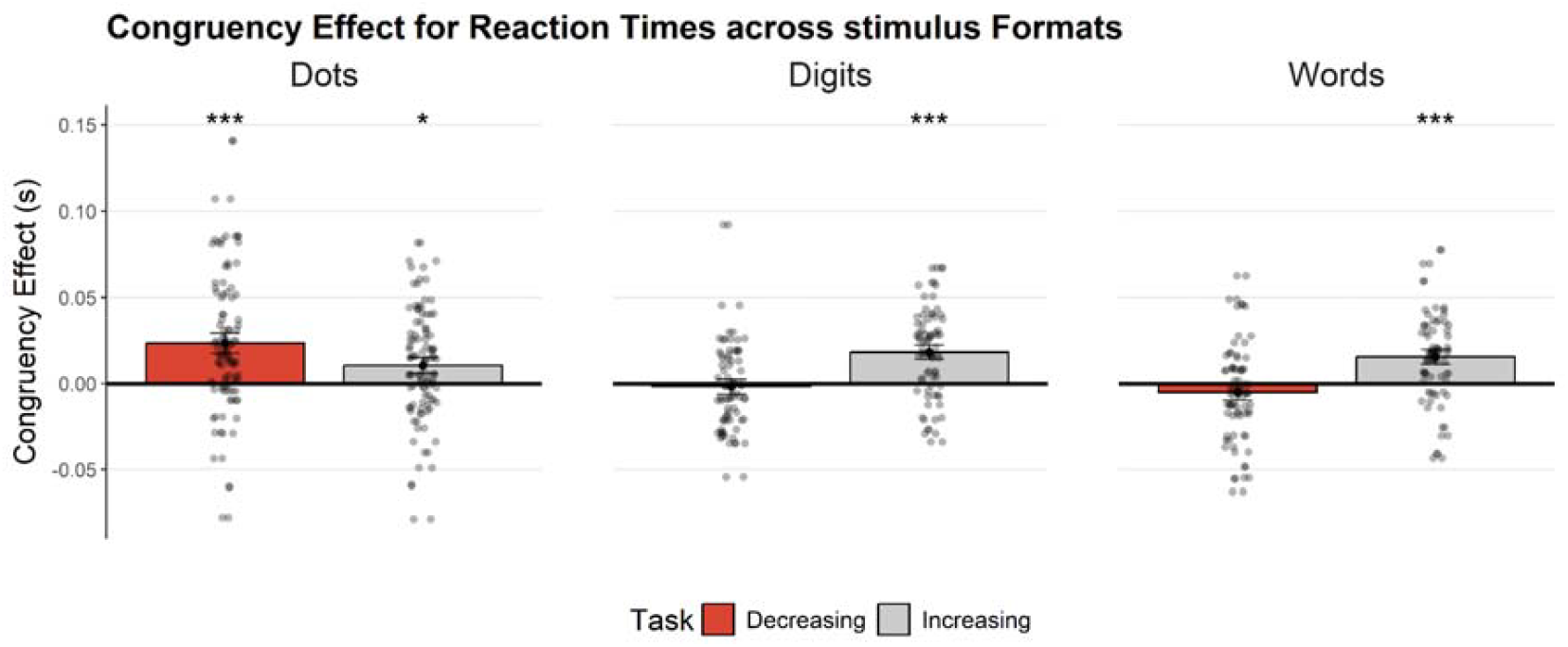
Congruency Effect for each Task and Format. Means and standard errors of the mean are reported. Statistical significance for Two-tailed one-sample Wilcoxon signed rank test against chance level (µ = 0) is reported as follows: * = pvalue < .05, *** = pvalue <.001.

In addition, we compared the current set of data with the Italian adults data collected in Eccher et al., 2025 to investigate differences in RTs in function of the different numerical comparisons presented in the two tasks. A 2×2 ANOVA investigating the effect of study differences (Study), and Task revealed a main effect of Study (F_(1,96)_ = 38.626, p_value_ < .0001, η^2^_p_ = 0.287) and Task (F_(1,96)_ = 11.114, p_value_ < .001, η^2^_p_ = 0.104). Overall, shorter RTs were collected in Eccher et al., 2025 (mean = 0.450 s, SEM = 0.011) compared to the current work (mean = 0.560 s, SEM = 0.009), and, overall, in the increasing task (mean = 0.495 s, SEM = 0.011) compared to the decreasing task (mean = 0.520 s, SEM = 0.011). See Supplementary Materials for additional explorative analysis of study differences between each task (Figure S2 and Table S1).

## Discussion

In our previous work (Eccher et al., 2025), we demonstrated that when using an implicit task, SNAs emerge independently from reading/writing habits. However, while we found that small numerosities were responded to more efficiently when they were presented on the left hemifield (thus emerging in our “decreasing”, or “press when smaller” task), we did not detect a convincing symmetrical facilitation for large numerosities presented on the right visual hemifield (that should, but did not, emerge in our “increasing”, or “press when more” task). We hypothesised that this asymmetry might have been determined by either of two factors (or by their combination): first, non symbolic numerosity tends to be better represented in the right hemisphere, masking the congruency SNA effect to emerge when stimuli were presented in the right hemifield; second, increasing changes tend to be faster and more automatically perceived, thus potentially masking congruency SNA effect to emerge when subjects performed this task, due to their high speed (Ben-Meir et al., 2012; Ganor-Stern, 2015; Müller & Schwarz, 2008).

In the current study, to test these hypotheses, we modified the original paradigm in two ways: first, in order to verify the idea that some of the asymmetry observed in the previous work might have been due to right hemispheric superiority in non-symbolic number processing, we presented dot patterns (similarly to the original Eccher et al. study) as well as digits and words, that are symbolic stimuli that are known to elicit a stronger left-hemispheric lateralization. Second, to reduce the automaticity in the detection of increasing numerosities and thus make the task more difficult, we introduced different and harder-to-detect trials, spanning 4 levels of increasing distance from the reference stimulus.

With respect to the hypothesis related to stimulus format (symbolic -digits and number words-*vs*. non-symbolic -dot patterns-), the results are in line with our predictions: for both digits and number words we observe a faster response when increasing numbers are presented on the right *vs*. the left, and no facilitatory response when decreasing numbers are presented on the left, thus a Congruency Effect only for the increasing task (Figure 3). At the same time, we confirmed and extended the results of Eccher et al. (2025): when non-symbolic stimuli were used, we observed a stronger Congruency Effect in the decreasing task. These results are in line with the idea that hemispheric specialisation for one or the other numerical stimulus format (Sokolowski et al., 2021) interferes with the ability to detect SNA using this paradigm: Symbolic stimuli preferentially activate the left hemisphere, leading to better processing of stimuli appearing on the right hemifield, thus masking the SNA effect in the decreasing condition. Importantly, this left hemisphere dominance alone cannot explain the effect observed, as notably, no reversed Congruency Effect was found for the decreasing task (i.e., right hemispace). Rather, as it happened in Eccher et al., 2025, these findings seem to be the result of the combined effect of ANS hemispheric specialisation and SNA.

Regarding our second hypothesis, that of the effect of task difficulty, we reasoned that introducing more demanding numerical comparisons would slow response times and, in turn, allow stronger SNA effects to emerge. Consistent with this hypothesis, analyses of raw reaction times confirmed that the tasks used in the present study were overall more difficult than those of the initial study. Specifically, the numerical comparisons used in this study (spanning from 0.75 to 0.94 ratios) were more difficult than those in the original paradigm (0.3 ratio), resulting in overall slower RTs. Critically, while in Eccher et al., 2025 the faster responses in the increasing task could have masked the SNA effect, here, with slower overall RTs we were able to find a Congruency Effect in both task conditions (although with a smaller effect size), likely due to higher task demands. Supporting this interpretation, previous research has shown that the strength of the SNARC effect is positively correlated with reaction times, with slower responses yielding stronger spatial-numerical associations (Cipora et al., 2019; Gevers et al., 2006). However, our findings reveal a more complex relationship between reaction times and the SNA effect. Notably, in the Arabic digit task, participants exhibited a robust SNA effect in the increasing task, despite relatively shorter reaction times.

Together, these results suggest that task demands, at least as indexed by reaction time duration, do not straightforwardly predict the presence or strength of the SNA effect. Rather than reflecting a linear relationship, the association between task demands, reaction times, and SNA may follow a non-linear pattern. One possibility is a reversed U-shaped relationship, akin to that described by the Yerkes–Dodson law (1908), whereby spatial– numerical associations are strongest within an optimal temporal window of response execution. In line with this hypothesis, future studies should explicitly investigate the dependency between response times and the SNA effect by systematically manipulating task demands, for example through variations in stimulus onset latency or the use of response-delay paradigms. Importantly, such approaches should also assess whether these dynamics differ across stimulus formats, as the present data suggest that reaction time variations do not modulate SNA in the same way for symbolic and non-symbolic stimuli.

In conclusion, our results provide further evidence that SNA is a complex phenomenon shaped by both biological and cultural factors. We show that left-to-right spatial–numerical associations in adult humans emerge differently depending on stimulus format and response time, suggesting that specific numerical notations may bias attentional orienting toward one hemispace over the other. While these findings offer new insights into the conditions under which SNA emerges, further research is needed to clarify how stimulus characteristics, task demands, and temporal dynamics interact, and to elucidate the neural mechanisms underlying these effects.

## Data and code availability

The data and the R code used to conduct the statistical analysis and to generate the plot figures are available at OSF repository

https://osf.io/ts8xe/overview?view_only=f03549b4ddb64c9e839f05eda90b5b58

## Acknowledgements

This work was supported by funding from the European Research Council under the European Union’s Horizon 2020 research and innovation program (Grant Agreement 833504 SPANUMBRA) to G.V. and under the ERC Proof of Concept programme (Grant Agreement 1117 No. 101189208, TaGaDeDys) to G.V.

## Supplementary Materials

**Figure S1.**
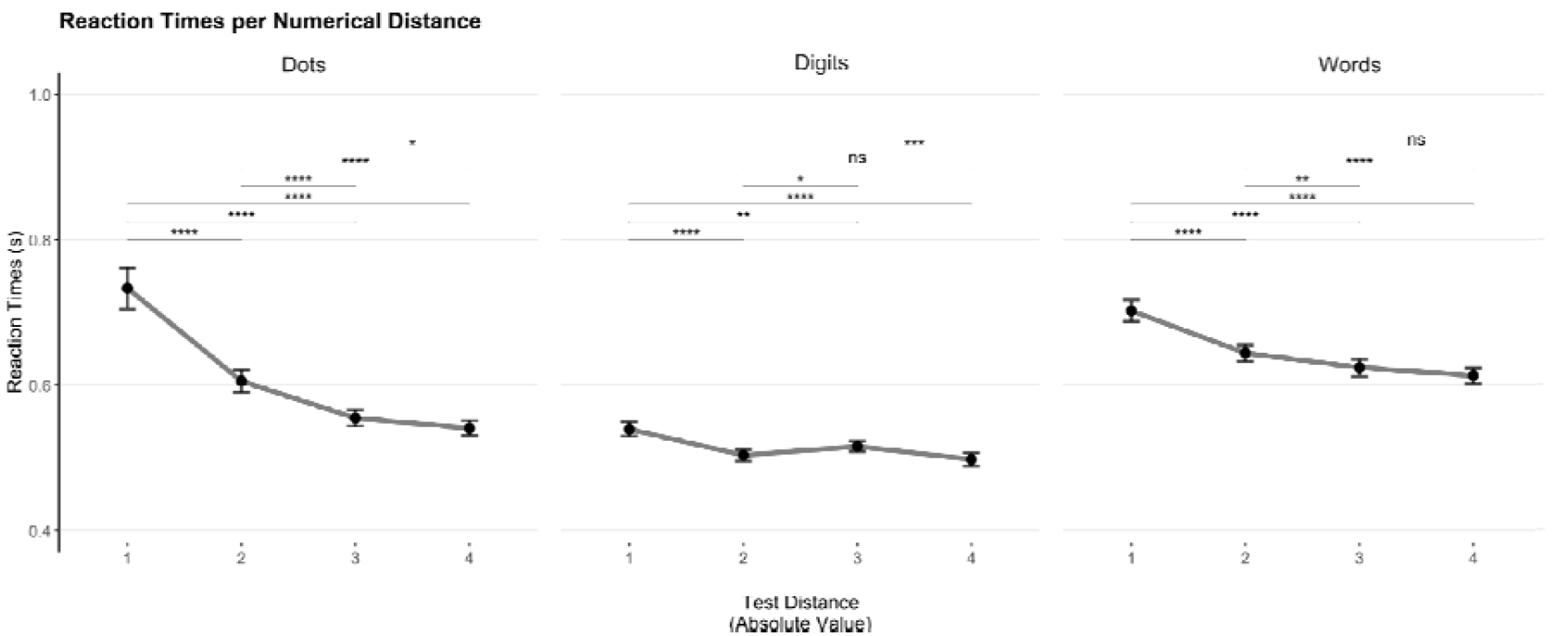
Reaction Times for numerical distance between reference and stimulus test, in absolute value, for each Format. In the graph the means and the standard error of the mean are reported by the black dots and error bars. Two-tailed two-sample Wilcoxon signed-rank test statistic, * = pvalue <.05, ** = p_value_ <.01 and *** = p_value_ <.001

**Figure S2.**
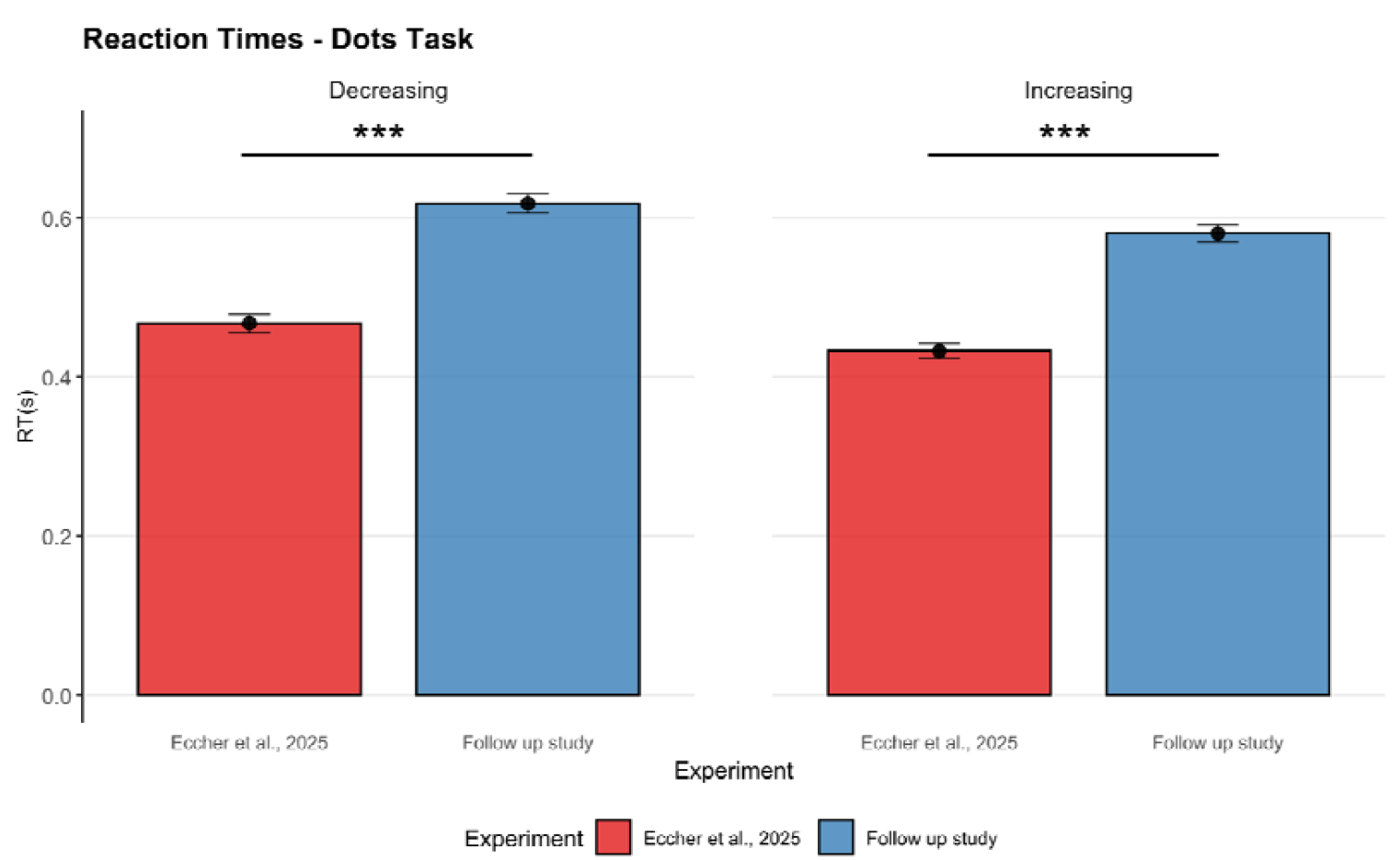
Reaction Times in Decreasing (left) and Increasing (right) tasks collected in Eccher et al., (red) and in the current work (blue). In the graph the means and the standard error of the mean are reported by the black dots and error bars. Two-tailed two-sample Wilcoxon rank-sum test statistic, *** = pvalue <0.001

**Table S1.**
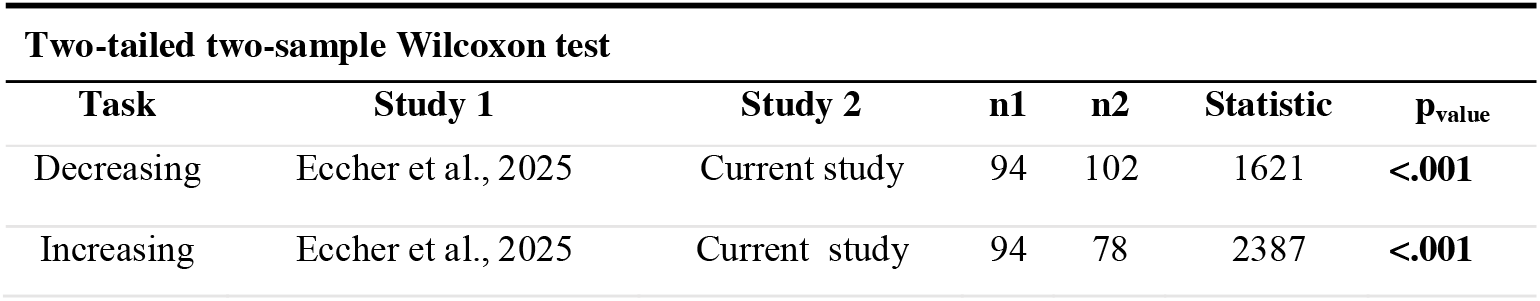
Statistical comparisons between Reaction Times in the two studies for each Task. Explorative analysis for differences between studies in each task. Significant pvalues for alpha = .05 in bold.

| Two-tailed two-sample Wilcoxon test |  |  |  |  |  |  |
| --- | --- | --- | --- | --- | --- | --- |
| Task | Study 1 | Study 2 | n1 | n2 | Statistic | Pvalue |
| Decreasing | Eccher et al., 2025 | Current study | 94 | 102 | 1621 | <.001 |
| Increasing | Eccher et al., 2025 | Current study | 94 | 78 | 2387 | <.001 |

